# Spatiotemporal sensitivity of embryonic heart specification to FGFR signaling in *Drosophila*

**DOI:** 10.1101/2020.11.16.384123

**Authors:** V. Yadav, N. Tolwinski, T. E. Saunders

## Abstract

Development of the *Drosophila* embryonic mesoderm is controlled through both internal and external inputs to the mesoderm. One such factor is Heartless (Htl), a Fibroblast Growth Factor Receptor (FGFR) expressed in the mesoderm. Htl is involved in shaping the mesoderm at both early and later stages during embryogenesis. How Htl expression levels and timing of signaling affect mesoderm morphogenesis after spreading remains elusive. We have developed an optogenetic tool (Opto-htl) to control the activation of Htl signaling with precise spatiotemporal resolution *in vivo*. We find that the embryo is most sensitive to Htl over-activation within a developmental window of ~4 hours ranging from late stage 10 until early stage 13, which corresponds to early stages of heart morphogenesis. Opto-htl restores heart cells in *htl* mutants upon light activation, independent of its role in early mesoderm shaping events. We also successfully generated spatially distinct regions of Htl activity in the mesoderm using light-sheet microscopy. The developing tissue was unable to correct for the ensuing asymmetries in cell fate. Overall, Opto-htl is a powerful tool for studying spatiotemporal regulation of Htl signaling during embryogenesis.

## INTRODUCTION

Morphogenetic events are tightly controlled in space and time (Wartlick *et al*., 2011; Ebisuya & Briscoe, 2018; Huang & Saunders, 2020). Throughout development, cells propagate, migrate, and differentiate to form specialised structures in a highly regulated manner. Such self-organisation within the embryo is dictated by numerous signaling events, some of which are highly conserved across organisms (Wangler *et al*., 2015; Beira & Paro, 2016; Li & Elowitz, 2019). Many of these signaling pathways interact with each other, generating complex network interactions and are interpreted differently by cells based on their competence to respond (Perrimon *et al*., 2012; Sagner & Briscoe, 2017; Yin *et al*., 2018; Li & Elowitz, 2019). These signaling networks are regulated in both space and time throughout morphogenesis to ensure that development is robust and reproducible (Amourda *et al*., 2017; Huang *et al*., 2017; Zinner *et al*., 2020).

A crucial signaling pathway in regulating cell behaviour during development is the highly conserved Fibroblast Growth Factor Receptor (FGFR) pathway (Sopko & Perrimon, 2013). FGFRs are transmembrane proteins that belong to the receptor tyrosine kinase family. Ligand-binding leads to FGFR homo-dimerisation, initiating trans-phosphorylation events which ultimately activates transcription of several target genes (Fantl *et al*.,1993). This transcriptional response regulates different cellular responses, such as changes in cell morphology, proliferation, adhesion, migration and differentiation (Spivak-Kroizman *et al*., 1994; Mohammadi *et al*., 1996; Thisse & Thisse, 2005; Teven *et al*., 2014; Mele & Johnson, 2020). Due to their involvement in key cellular processes, FGFRs when mutated can promote cancer development and progression (Turner & Grose, 2010; Wesche *et al*., 2011; Presta *et al*., 2017).

There are two known FGFRs in *Drosophila* that control distinct developmental processes, *heartless (htl)* and *breathless (btl)*. These bind three ligands in total - Pyramus, Thisbe and Branchless (Stathopoulos *et al*., 2004; Kadam *et al*., 2009). This compares with four known FGFRs and at least 22 associated ligands in humans that bind each other in several different combinations (Ornitz & Itoh, 2015). The small number of receptor-ligand combinations makes *Drosophila* an excellent system to explore the basic interactions underlying FGFR action. In *Drosophila, htl* expression is important for proper development of several mesoderm derived tissues, including the heart and muscles (Beiman *et al*., 1996; Gisselbrecht *et al*., 1996; Shishido *et al*., 1997). *btl* expression is required for proper morphogenesis of the trachea (Klambt *et al*, 1992; Reichman-Fried *et al*., 1995). Both Htl and Btl are essential in driving proper migration of mesodermal, glial, and tracheal cells (Muha & Müller., 2013). For instance, Htl plays a role in the spreading of the mesoderm over the ectoderm to form a monolayer during early stages of embryogenesis (Wilson *et al*., 2004; Wilson *et al*., 2005; McMahon *et al*., 2010; Bae *et al*., 2012). Uniform spreading of the mesoderm is crucial for proper cell-fate specification of different cell types within the mesoderm at later stages (Stathopoulos *et al*., 2004; Kadam *et al*., 2009). In *htl* mutants, mesoderm cells fail to undergo proper spreading and form irregular and multilayer arrangements. This lack of structure prevents mesodermal cells from receiving precise spatial cues from the ectoderm. Later in development, Htl is also involved in the specification of different cell types derived from the mesoderm (Michelson *et al*., 1998). *htl* null mutants lack precursors of pericardial and heart cells, have defects in visceral mesoderm, and show reduced, irregular muscle patterns (Beiman *et al*., 1996; Stathopoulos *et al*., 2004; Kadam *et al*., 2009).

The role of FGFR in cell fate specification has been extensively studied (Carmena *et al*., 1998; Ciruna & Rossant., 2001; Mandal *et al*., 2004). While previous work has provided detailed insights into how Htl controls the movement of mesodermal cells during the spreading phase (McMahon *et al*., 2008), the *in vivo* dynamics of Htl action within the developing mesoderm remain elusive after the initial stages of spreading. Genetic perturbations of *htl* offer only a limited exploration of the spatiotemporal range of Htl activity. In recent years, the use of optogenetics to tune signaling pathway responses has become a powerful tool *in vivo* (Huang *et al*., 2017; Kaur *et al*., 2017; Viswanathan *et al*., 2019; Johnson *et al*., 2020). Optogenetic approaches enable precise spatiotemporal tuning of target activity, enabling *in vivo* signaling dynamics to be dissected.

Here, we utilised an optogenetic tool (termed Opto-htl) to activate Htl signaling in a spatiotemporally controlled manner during *Drosophila* embryo development. Upon illumination with 488nm light, Opto-htl functions as a constitutively active receptor, capable of activating downstream factors of the FGFR pathway, such as the extracellular signal regulated kinase (Erk). Opto-htl restored a significant number of heart cells within a *htl* mutant upon light activation, though it did not fully rescue the mutant phenotype. Constitutive activation in the mesoderm of the wild-type led to several developmental defects, the severity of which varied with changes in light intensity, timing, and spatial organisation of the light exposure. We identified a time window of sensitivity to FGFR over-activation (stage 10 till late stage 12 of embryogenesis), illumination during which was both necessary and sufficient to induce the phenotypic defects. Together, these results demonstrate sensitivity of the Htl-dependent processes (particularly heart formation) to over-activation of Htl.

## RESULTS

### Opto-htl can stimulate FGFR activity

To generate an *in vivo* optogenetic tool for FGFR activation, we utilised Cryptochrome2 (CRY2), a light-interacting molecule that undergoes dimerisation upon exposure to 488nm blue light (Yu *et al*., 2009). The cytoplasmic domain of *htl* was fused with CRY2-mCherry and the resulting fusion protein (termed Opto-htl) was anchored to the membrane by a myristoylation (myr) signal sequence (Kim *et al*., 2014). Light exposure induces dimerisation of CRY2, bringing two receptor molecules together and triggering a phosphorylation cascade, which should lead to ligand-independent activation of target genes downstream of the receptor (Fig. 1A).

**Fig. 1.**
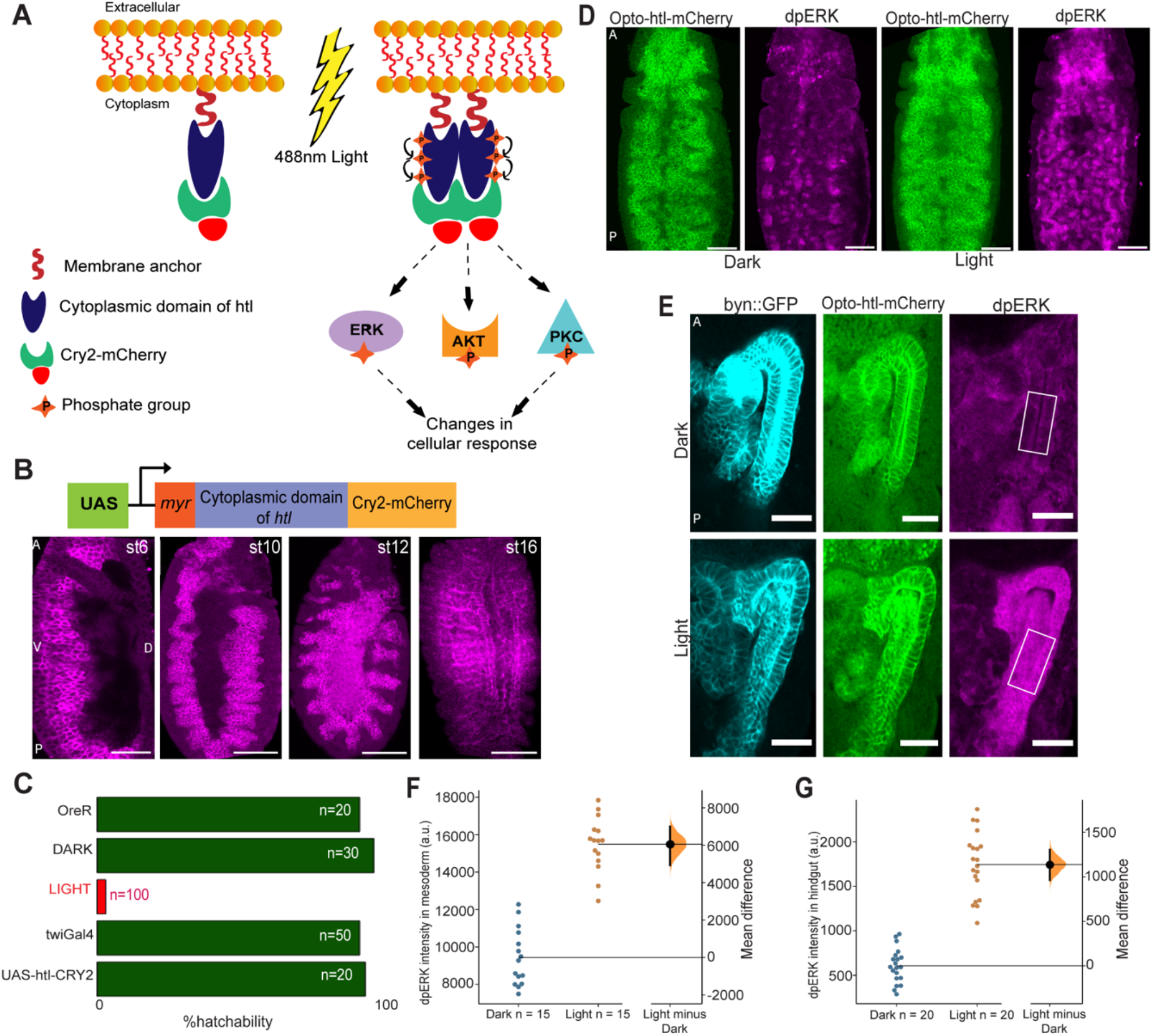
Opto-htl expression and activation in different tissues. A) Schematic showing design and activation of Opto-htl. Exposure to 488nm light induces CRY2 clustering leading to activation of the (intracellular) membrane-bound receptor and eventually generating a cellular response via phosphorylation of various downstream target molecules. B) Twi-Gal4>Opto-htl embryos stained with anti-mCherry antibody showing the expression of the construct at various stages. C) Hatchability assay for different genetic conditions. OreR = OregonR, DARK = Twi-Gal4>Opto-htl embryos kept under illumination with amber paper to block 488nm wavelengths throughout development, LIGHT = Twi-Gal4>Opto-htl embryos kept under illumination throughout development. Twi-Gal4 embryos represent the Gal4 driver alone. UAS-htl-Cry2-mCherry represents Opto-htl embryos with no Gal4 driver. D) Twi-Gal4>Opto-htl embryos fixed and stained at late stage 10/early stage 11 for pErk under dark and light conditions. E) Byn-GAL4>Opto-htl embryos fixed and stained at stage 16 for pErk under dark and light conditions (scale bar 25μm). F) pErk intensity differences in the mesoderm at late stage 10 between Twi-Gal4>Opto-htl embryos kept under dark versus light. G) pErk intensity differences in the hindgut between Byn-GAL4>Opto-htl embryos under dark and light conditions in stage 16 measured in the region shown in (E). Scale bar = 50μm unless stated otherwise. A = Anterior, P = Posterior, D = Dorsal, V = Ventral view. In (F,G), the black bar represents the 95% confidence interval with the Bootstrap distribution shown in orange (see Ho *et al*., 2019 for details).

We used Twi-Gal4 to drive Opto-htl (Twi-Gal4>UAS-htl-CRY2-mCherry) (Fig. 1B, top) in the mesoderm. We validated that our construct was expressed in the mesoderm and mesoderm-derived tissues at different stages throughout embryogenesis (Fig. 1B). Live imaging showed that expression levels remained low until stage 10 but subsequently increased and were maintained in the mesoderm throughout embryogenesis (Movie S1). Twi-Gal4>Opto-htl embryos maintained in the dark throughout embryogenesis hatched with similar frequency to wild-type (OreR) embryos (Movie S3). However, upon constant light exposure (Methods) from stage 5 onwards, there was a significant drop in hatchability and 97% of embryos died before hatching (Fig. 1C). These embryos developed up until the later stages of embryogenesis before dying unhatched (Movie S3).

Next, we tested the constitutive activation of the FGFR pathway via Opto-htl. We used the phosphorylation of Erk as a readout for the MAPK/Erk pathway downstream of FGFR (Gabay *et al*., 1997). Erk activation to doubly phosphorylated Erk (dpErk) leads to transcription of several target genes that regulate growth and proliferation (Lavoie *et al*., 2020). We tested for differences in the levels of dpErk in Opto-htl embryos kept under dark and light conditions. We illuminated Twi-Gal4>Opto-htl embryos up until the end of germ band elongation (late stage 10 / early stage 11), then fixed and stained them with anti-dpErk antibody alongside embryos kept in the dark. We observed an increase in dpErk levels in embryos exposed to light as compared to the ones fixed and stained at a similar stage in the dark (Fig. 1D). We also tested Erk activation in the hindgut epithelium (ectoderm-derived) which does not express high levels of dpErk in wild-type conditions. We used Byn-Gal4 to express Opto-htl in the hindgut epithelium and again tested for dpErk levels under dark and light conditions (Fig. 1E). Comparing the dpErk intensity for dark and light conditions in both tissues, dpErk levels were significantly increased upon light activation (Fig. 1F-G). We conclude that Opto-htl can be used to induce Erk activation *in vivo* upon illumination.

### Opto-htl can induce heart cells in a *heartless* mutant

To further test the functionality of Opto-htl, we expressed it against a null *htl* mutant background to explore the extent of rescue of phenotypes at different stages upon light illumination. Homozygous mutants for *htl* undergo improper spreading of mesoderm cells at stage 10 forming irregular and multilayer arrangements (Stathopoulos *et al*., 2004), lack pericardial precursors, a proper muscle structure, pattern and, as the name suggests, fail to form a proper heart (Beiman *et al*., 1996) (Fig. 2A). A previous tool to induce *htl* over-expression, htl-λ (Michelson *et al*., 1998; Wilson *et al*., 2005), was able to partially rescue the *htl* mutant, with some Eve-positive cells (precursors for pericardial and dorsal muscles) in the mesoderm restored.

**Fig. 2.**
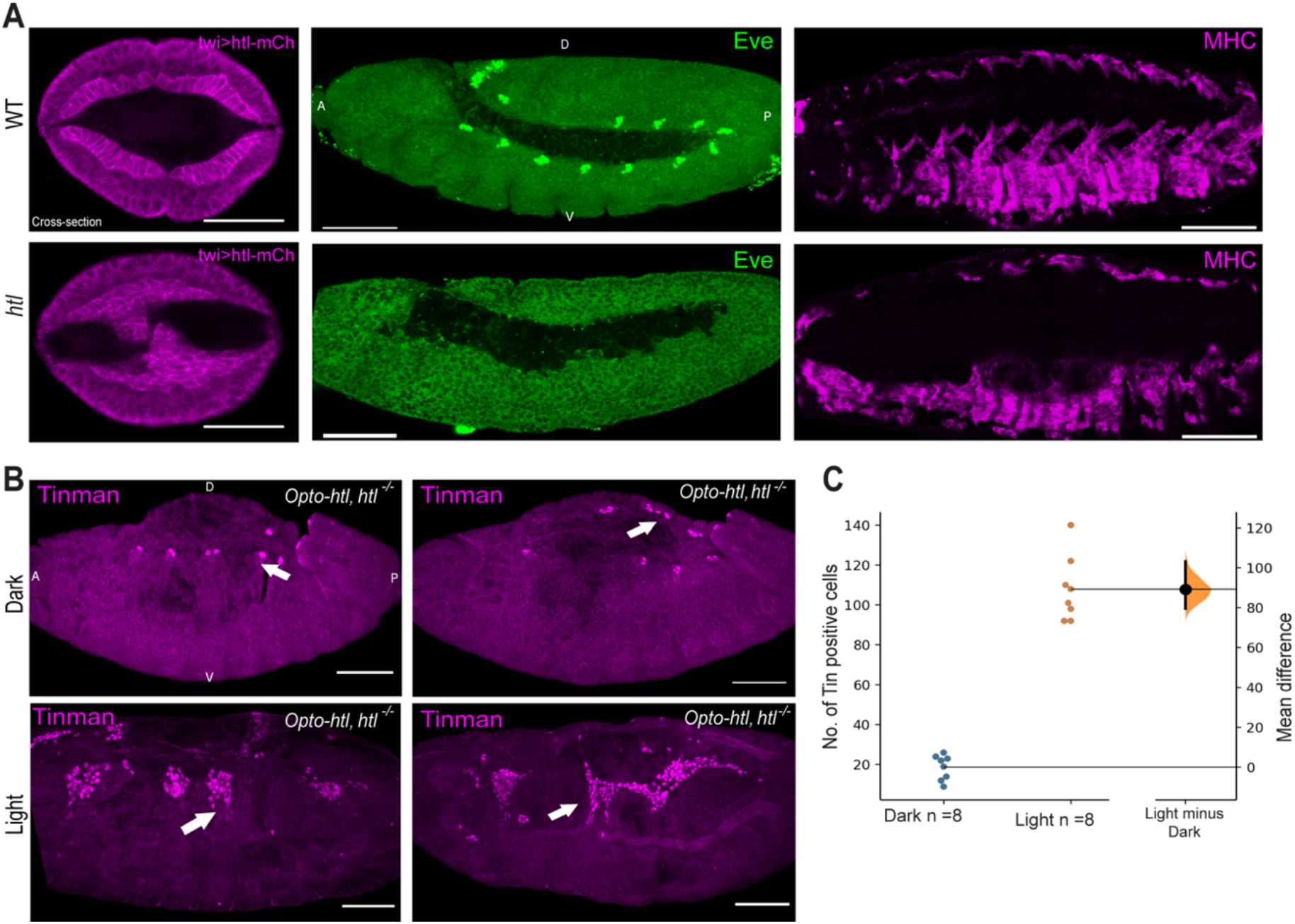
Rescue of *htl* mutant using Opto-htl. A) Comparison between wild-type and *htl* homozygous mutant embryos at different stages: Mesoderm spreading (stage 10), Eve-positive pericardial precursor cells (stage 11) and muscles marked by Myosin staining (stage 15). B) Twi-Gal4>Opto-htl is expressed against *htl* null background and embryos illuminated and stained with Tin antibody. Two homozygous mutants are compared under dark and light conditions for Tin-positive cells. C) Quantification of the number of Tin-positive cells in dark and light conditions for embryos as described in (B). Scale bar = 50μm. A = Anterior, P = Posterior, D = Dorsal, V = Ventral view. In (C), the black bar represents the 95% confidence interval with the Bootstrap distribution shown in orange (see Ho *et al*., 2019 for details).

We tested for rescue of heart cells at stage 16 in homozygous *htl* mutants expressing Twi-Gal4>Opto-htl with and without light activation. Tinman (Tin) is a transcription factor required for the specification of all heart cells and a subset of muscles (Bodmer, 1993). At stage 15/16, Tin is expressed in a major portion of the cardioblasts and pericardial cells (Alvarez et al., 2003). We found that homozygous embryos maintained in light showed restoration of Tin-positive heart cells at stage 16 when compared to embryos kept in the dark (Fig. 2B-C). In comparison, in wild-type embryos, there are around 104-Tin-positive cells in hemisegments A2-A8 (52 cardioblasts and pericardial cells each) at late stage 15/16 (Alvarez et al., 2003). Although Tin-positive cells were greatly increased under light activation of Opto-htl in *htl* null embryos, there was substantial variation in their spatial arrangement between embryos. The Tin-positive cells were located near the embryonic midline, but they failed to form a coherent heart structure. This is likely a consequence of embryos failing to undergo uniform mesoderm spreading over the ectoderm early on.

Opto-htl showed no rescue of phenotypes during early stages of development (Fig. S1). Since Opto-htl expression levels are generally low during early stages (Movie S1), it is possible that rescue during this stage requires higher levels of FGFR signaling than we currently achieve. These results are consistent with previous findings that Htl-dependent cell fate decisions in the mesoderm after stage 10 are decoupled from its role in mesoderm spreading (Michelson *et al*., 1998), though formation of robust organs requires synergy between both processes.

### Activation of Opto-htl induces ectopic Tin-positive cardioblasts

Light activation of Opto-htl in the wild type mesoderm interfered with embryonic development, leading to a significant increase in lethality. We focused on the effect of Opto-htl activation on the development of mesoderm-derived tissues, particularly the heart. The heart in wild-type and Opto-htl embryos kept under dark is composed of two rows of cells (Fig. 3A left) with a repeated pattern of four Tin-positive and two-Seven-up (Svp) positive cells (Ahmad, 2017; Vogler & Bodmer, 2015) (Fig. 3B, top). The cells comprising these two rows are specified on both lateral sides of the embryo by the end of stage 12 after which they start migrating towards each other and match in a highly precise fashion (Reim & Frasch., 2010, Zhang *et al*., 2018; Zhang *et al*., 2020) (Fig. 3B top, Movie S4). Upon illumination, the stage 16 heart in Twi-Gal4>Opto-htl embryos showed a significant increase in the number of cardioblasts (Fig. 3A right and Fig. 3B bottom). There was an increase in the number of cardioblasts per hemisegment, but with substantial variation between embryos (Table 1). The clusters still had Svp-positive cells appearing in doublets, identifiable by reduced Fasciclin-III (Fas3) expression and morphology (Zhang *et al*., 2018) (asterisk in Fig. 3B) as in wild-type embryos, but they no longer matched with their contralateral partners. In some cases, the heart structure even becomes discontinuous (Movie S5). The observed asymmetries in the light-activated Opto-htl embryos suggested that the ectopic heart cells were generated asymmetrically between the contralateral sides of the embryos. Some embryos even developed multiple branches emanating from what appeared to be the primary heart vessel (arrowhead Fig. 3B).

**Fig. 3.**
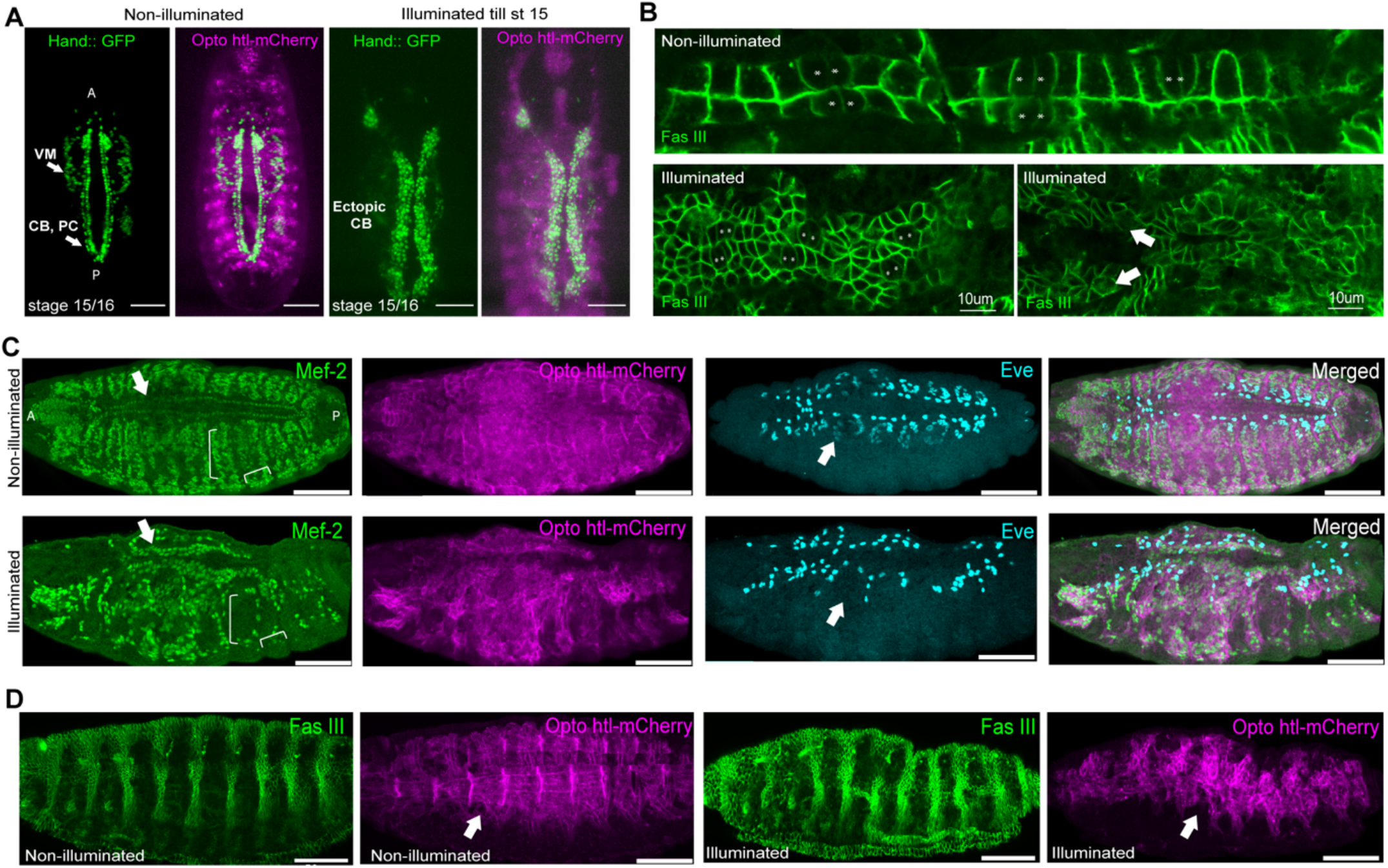
Mesoderm specific defects in Twi-GAL4> UAS - htl-CRY2-mCherry embryos kept under constant light. (A) Twi-GAL4>Hand::GFP;Opto-htl embryos kept under dark and illuminated conditions, imaged using a light-sheet at stage 15 (Opto-htl signal in magenta). (B) Twi-GAL4>Opto-htl embryos stained with Fas3 and imaged at stage 16 under dark conditions (top) and when illuminated by constant 488nm light (bottom). Svp-positive cells, marked with an asterisk, are identified by lower Fas3 expression. Arrows denote branching of the heart structure. (C) Twi-GAL4>Opto-htl embryos stained with Mef-2 and Eve antibodies at stage 16 under dark and illumination conditions. Arrows correspond to phenotypes described in the text. (D) Muscle structure in Twi-GAL4>Opto-htl embryos kept under dark and light conditions visualised using Opto-htl-mCherry. Scale bar = 50μm unless stated otherwise.

**Table 1:** Number of Mef2-positive cardioblasts, Eve-positive pericardial cells and DA1 muscles in OreR, Twi-Gal4>Opto-htl (dark) and Twi-Gal4>Opto-htl (illuminated) embryos. Cells were counted at stage 15/16 from hemisegments A3-A4.

| Condition/Genotype | Mef2-CB | Eve-PC | DA1 |
| --- | --- | --- | --- |
| OreR<br>(n=12) | 12 | 4 | 2 |
| DARK-twi>Opto-htl<br>(n=10) | 12 | 4 | 2 |
| Light-twi>Opto-htl<br>(n=10) | 29 | 6 | 0 |
|  | 32 | 7 | 0 |
|  | 41 | 9 | 0 |
|  | 26 | 3 | 0 |
|  | 20 | 6 | 0 |
|  | 31 | 7 | 0 |
|  | 30 | 6 | 0 |
|  | 27 | 7 | 0 |
|  | 21 | 3 | 0 |
|  | 23 | 4 | 0 |

### Activation of Opto-htl disrupts mesoderm-derived muscle formation

In wild-type embryos, the dorsal muscles - DA1 muscles - are initially flower like structures formed from a cluster of Eve-positive cells around both sides of the heart (Han & Bodmer., 2003; Schwarz *et al*., 2018). Upon staining with Eve antibody, we observed that the DA1 muscle precursors are completely missing from Opto-htl embryos kept under light (Fig. 3C, Eve). These embryos also showed a reduced number of muscle-forming Mef2 positive cells (Fig. 3C, brackets) (Mef-2 is a key transcription factor that marks and directs proper specification of the heart and muscle cell types (Bour *et al*., 1995; Lilly *et al*., 1995)). We next visualised the muscle patterns at stage 16 using the mCherry signal marking Opto-htl. In wild-type embryos and in Opto-htl embryos kept in dark, the somatic muscles were arranged and attached to the body wall in regular patterns (Zhou *et al*., 2019) (Fig. 3D). This pattern was disrupted in Opto-htl embryos kept under light, with the muscles appearing diffused and disrupted (Fig. 3D). Overall, we see that over-activation of Opto-htl leads to a loss of structural organisation within mesoderm-derived tissues.

### Opto-htl allows spatially precise activation of Htl signaling within the developing mesoderm

We used a light-sheet microscope to generate a spatially restricted region of Opto-htl activation within a developing embryo. Since the light-sheet illuminates and collects the signal from a single optical section at a time, we are able to partition a given embryo into illuminated and non-illuminated sections. (Kaur *et al*., 2017). A Twi-Gal4>hand::GFP; Opto-htl embryo was mounted such that the illumination plane was parallel to the long-axis of the embryo. To generate spatially heterogeneous activation of Opto-htl, we scanned the Opto-htl embryos with the 488nm laser to a depth of ~35um from the embryo surface on one lateral side, so as to illuminate the heart precursors on only one side of the embryo (Fig. 4A). At stage 16, we imaged the embryo from the dorsal side to image the heart and compared Hand::GFP signals from the illuminated and non-illuminated sides. We observe a striking difference between the number of heart cells on both sides, with the non-illuminated side resembling the dark condition phenotype while the illuminated side showed multiple ectopic cardioblasts (Fig. 4B, asterisk).

**Fig. 4.**
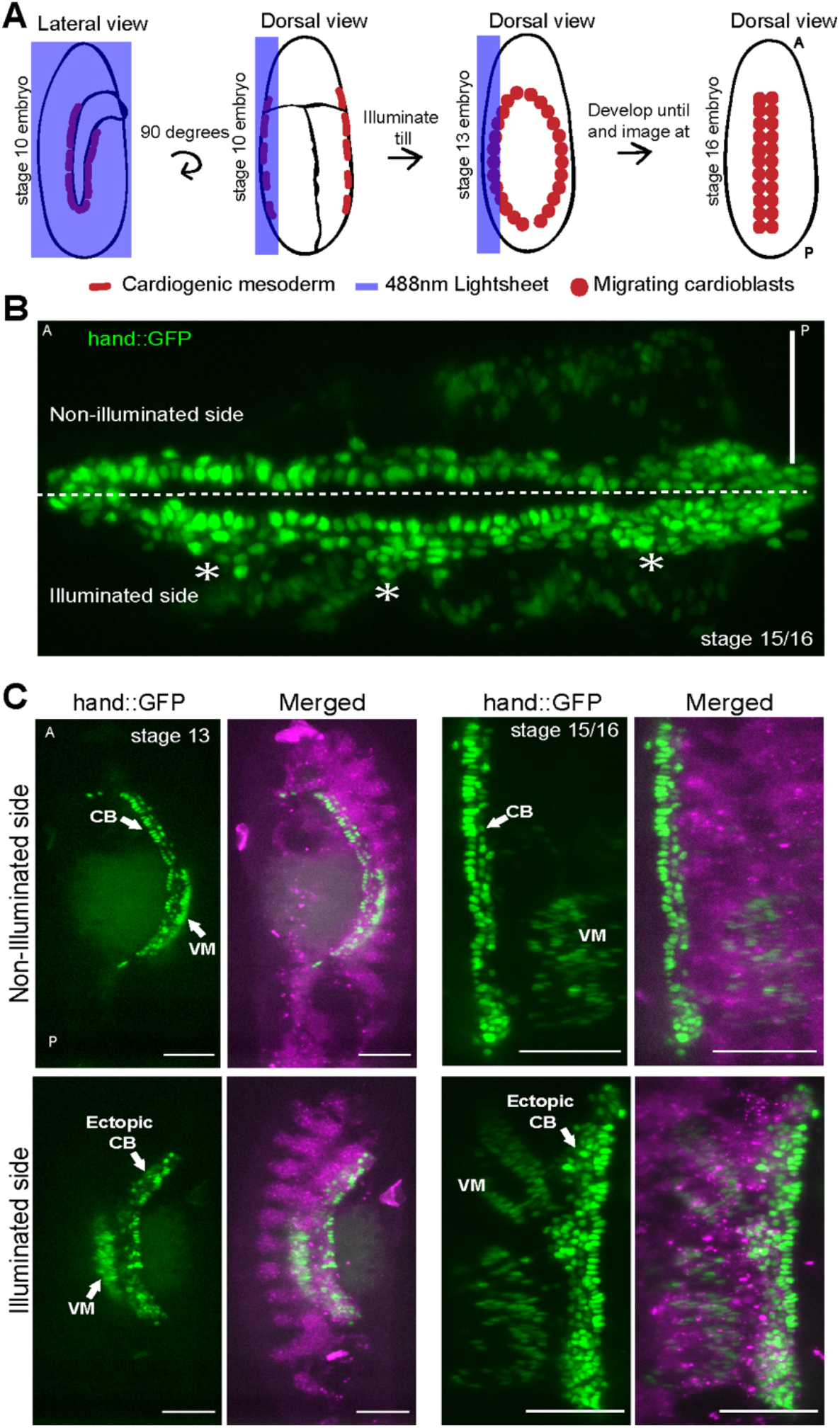
Spatial control of Opto-htl activation. A) Schematic of the imaging protocol for half-illumination of embryos on a light-sheet microscope. B) Dorsal view of a Twi-Gal4>hand::GFP;Opto-htl embryo illuminated as described in (A) and imaged at stage 16. C) Non-illuminated (top row) and illuminated (bottom row) sides of a Twi-Gal4>hand::GFP;Opto-htl embryo. Left panels, stage 13. Right panels, stage 15/16. VM= Visceral mesoderm, CB= Cardioblasts, PC= Pericardial cells. Scale bar = 50μm.

We observed ectopic cardioblasts in the illuminated side immediately after the end of illumination at stage 13 (Fig. 4C, left). By the end of stage 15 (Fig. 4C, right), the non-illuminated side resembled the wild type embryo in its arrangement and number of heart cells while the illuminated side had substantial ectopic cardioblasts arranged irregularly along the embryonic midline. These results suggest that the two lateral sides of the mesoderm develop independently from each other, and they cannot correct for defects in the formation of their contralateral partner which is consistent with the fact that the mesodermal cells do not mix extensively along the lateromedial axis and their spatial information with respect to one another is more or less conserved (Ortega & Hartenstein, 1985).

### Severity of Opto-htl induced phenotypes is dosage dependent

We observed the aforementioned phenotypic defects in Opto-htl embryos upon continuous and uniform illumination at a fixed intensity of 1mW (measured at the sample plane). Next, we varied the dosage of signaling via Opto-htl by illuminating Opto-htl embryos with different light intensities using a Nikon LED light base as the light source. We used light intensities at 2 mW, 1 mW, 0.25 mW, 0.1 mW, and 0.01 mW (measured using an intensity power meter set at the 488nm range) for illuminating Twi-Gal4>hand::GFP; Opto-htl embryos from stage 5 up until stage 15/16 and then imaged the heart in each embryo. Hand::GFP is expressed at stage 16 in all the heart cells and the adjacent visceral mesoderm. In Fig. 5A we depict the most and least defective heart images at each intensity, based on the number of ectopic cardioblasts. At low intensities (0.01mW), the least defective embryos closely resembled embryos maintained in the dark condition (Fig. 5B), with only a few ectopic cardioblasts. At 0.1mW, ectopic cardioblasts were more apparent in some embryos, though the overall structure still resembled the non-illuminated embryos. For intensities 0.25mW and above, the heart defects were marked, with major structural changes to the heart vessel (branching, inconsistent matching).

**Fig. 5.**
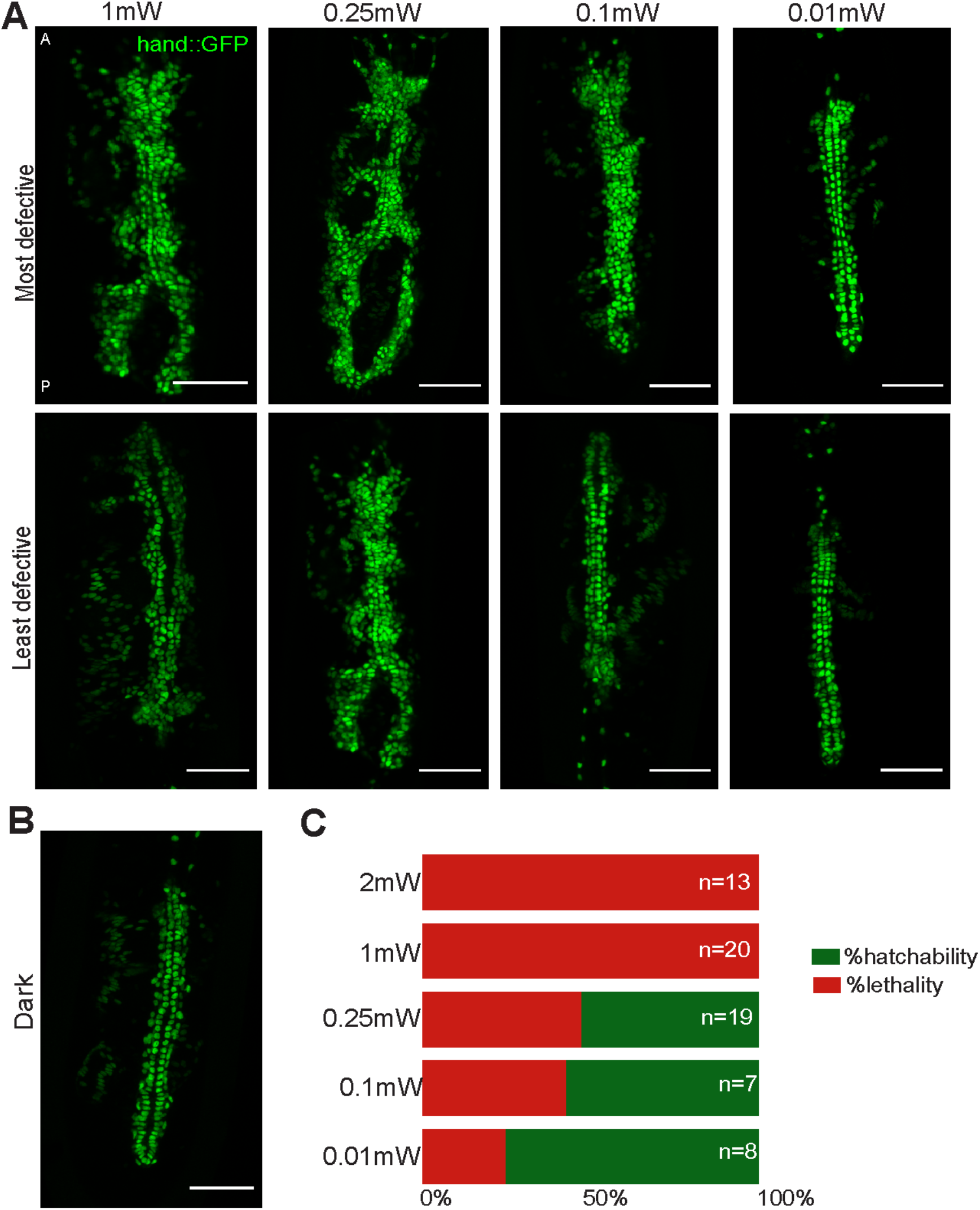
Opto-htl induced phenotype depends on light intensity. A) Representative images of the heart at stage 16 in Twi-Gal4>hand::GFP;Opto-htl embryos illuminated from stage 5 till stage 16, at different light intensities. B) Heart at stage 16 in a Twi-Gal4>hand::GFP;Opto-htl embryo kept in the dark through development. C) Survival rate of Twi-Gal4>Opto-htl embryos when illuminated at different intensities throughout development. Scale bar = 50μm.

After imaging the embryos at stage 16, we let them develop further and scored their hatchability. We found that increased intensity led to lower survival rates. At 0.1mW exposure, we observe a survival rate of around 57% as compared to a 100% lethality observed at intensity values of 1mW or higher (Fig. 5C). Despite significant ectopic cardioblasts, 52% of the embryos illuminated at 0.25mW still managed to hatch, possibly due to reduced severity of some other defects at lower intensities.

We compared these results with the other constitutively active form, htl-λ (Michelson *et al*., 1998; Wilson *et al*., 2005). htl-λ did not induce lethal phenotypic defects when expressed in the mesoderm. Tin and Eve patterns at stage 15 in Twi-Gal4>htl-λ embryos showed only small phenotypic variation from wild-type patterns (Fig. S3). The heart phenotype observed for htl-λ resembled that of Opto-htl embryos at lower intensity illuminations (<0.1mW). Of course, differences between the scales of action for the two constructs cannot be ruled out. At higher light activation, we see stronger phenotypes in the Opto-htl embryos as compared to htl-λ. Our results suggest that specification of the cardiac vessel within the mesoderm is highly sensitive to FGFR activity levels.

### Embryo development is sensitive to Opto-htl activation only during stages 10-12

In the early embryo, Htl plays an important role in regulating mesoderm spreading (Wilson *et al*., 2005; McMahon *et al*., 2010), but the dynamics of its activity later in development are less well known. We used Opto-htl to activate Htl signaling during distinct developmental time windows to disentangle its early and later effects on embryo development and to test when the developing mesoderm is most sensitive to Htl over-activation.

We illuminated Twi-Gal4>Opto-htl embryos during three separate time windows (Fig. 6A): (i) from stage 5 to stage 10 (~2.5 hours), during which the presumptive mesoderm forms the ventral furrow and undergoes spreading over the ectoderm to form a monolayer; (ii) from late stage 10 to late stage 12 (~4 hours), during which the specification of the presumptive mesoderm into different cell types occur; and (iii) from stage 13 up to stage 16 (~4 hours), during which cardioblasts migrate to the embryo midline and form the heart. We observe that nearly all embryos illuminated during windows (i) and (iii) hatch. However, almost all embryos illuminated between stages 10 and 12 fail to hatch (Fig. 6A). As previously stated, Opto-htl levels at early stages i.e. during time window (i), can only be detected by anti-mCherry antibody staining, suggesting low expression levels (Movie S1).

**Fig. 6.**
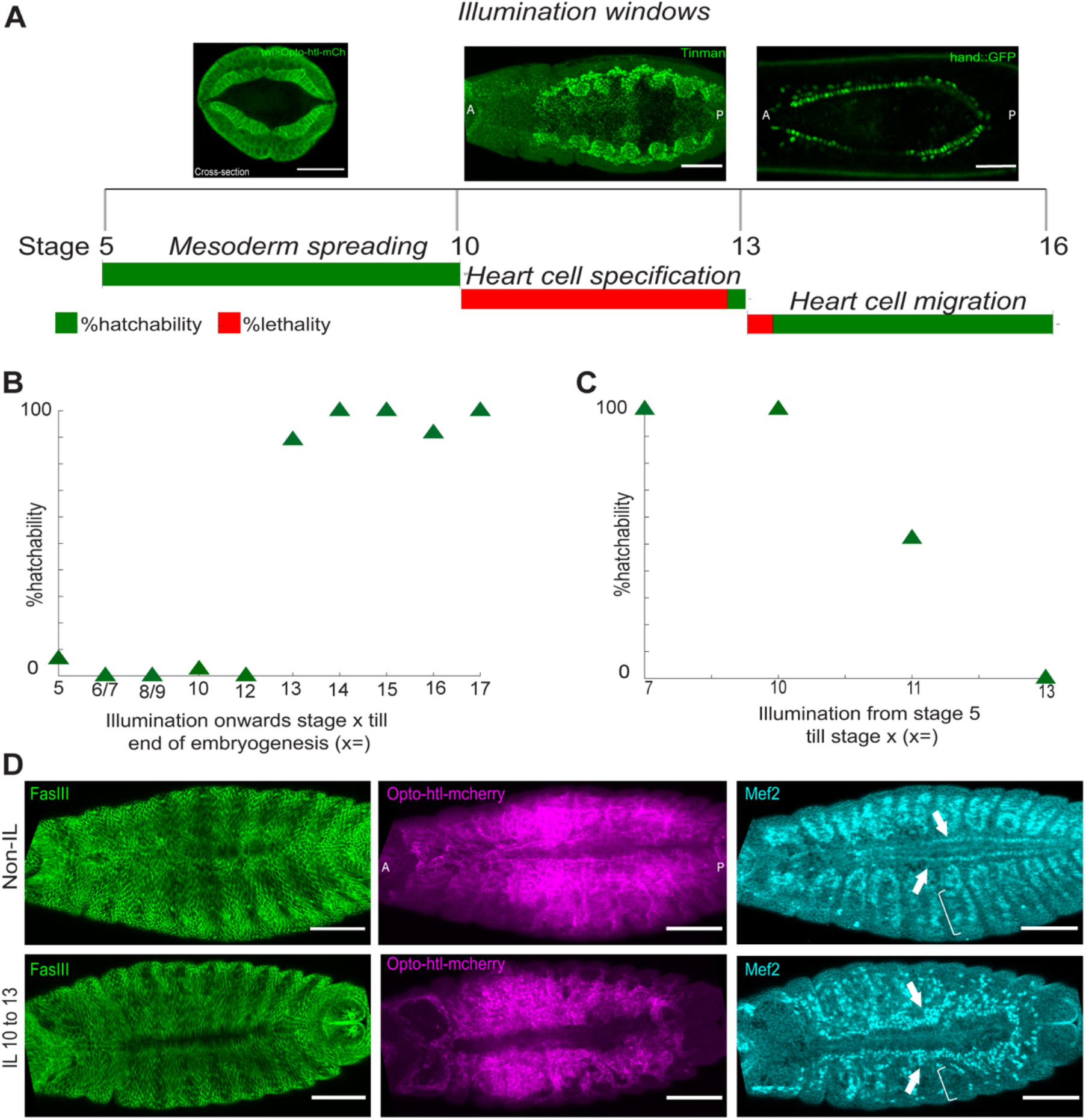
Temporal control of Opto-htl activation during development. A) Hatchability assay for Twi-GAL4>Opto-htl embryos illuminated during different time windows during development. (Top) Representative images of the embryo stage being illuminated. (Bottom) Hatching percentage, with the length of each bar representing 100%. B) Hatchability of Twi-GAL4>Opto-htl embryos when illuminated from the specified stage on the x-axis to the end of embryogenesis C) Hatchability of Twi-GAL4>Opto-htl embryos when illuminated from stage 5 until the specified stage on the x-axis (see Table 2A–B for embryo count) D) Twi-GAL4>Opto-htl embryos stained with Fas3, Mef2 and mCherry antibodies under different illumination conditions. Arrowheads point towards Mef-2 marked cardioblasts. Scale bars = 50μm.

We next performed illuminations during more fine-tuned developmental windows (Fig. 6B-C, Table 2A–B). We see a sharp transition in survivability around stage 12 where embryos illuminated prior to stage 12 show high lethality, and embryos illuminated only after stage 12 show high survivability (Fig. 6B). Performing the opposite light illumination protocol, embryos illuminated until stage 10 but then kept in the dark all hatched. Embryos illuminated until stage 11 had a survivability of around 50%. Developmental processes around stage 11-12 appear to be especially sensitive to Opto-htl activation. The specification of different cardiac progenitors occurs during this stage (Reim & Frasch., 2010). To determine what defects were induced by illumination of Opto-htl during time window (ii), we let embryos illuminated during this window continue to develop in the dark from stage 13 onwards and fixed them at stage 16. We observed that these embryos had ectopic Mef-2 positive cardioblasts (Fig. 6D, arrows), similar to Opto-htl embryos illuminated throughout development. Further, the Mef-2 positive muscle cells were also reduced as seen in the body wall muscles (Fig. 6D, brackets).

**Table 2A:**
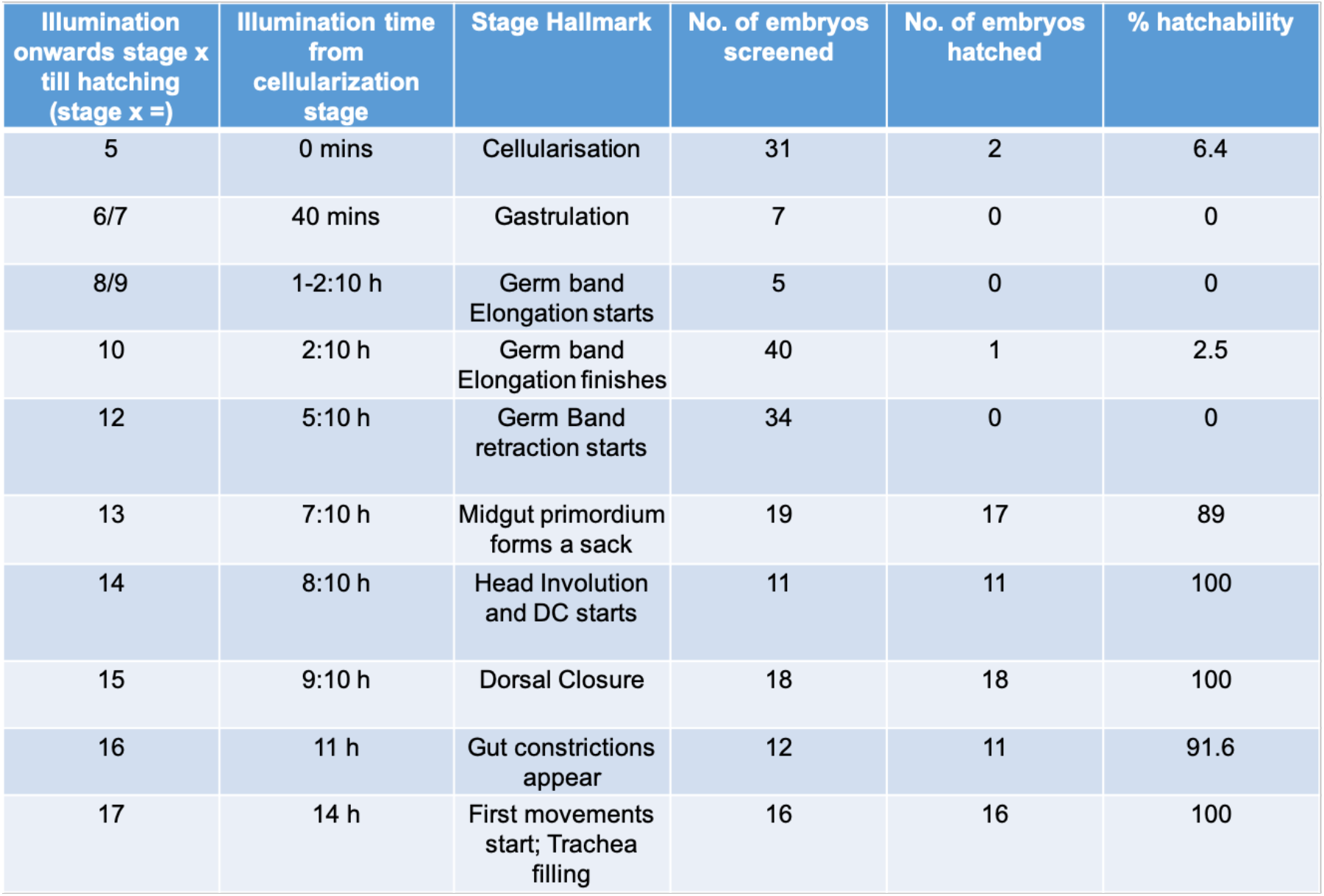
Hatchability of Twi-GAL4>Opto-htl embryos illuminated from specified stages until end of embryogenesis.

**Table 2B:**
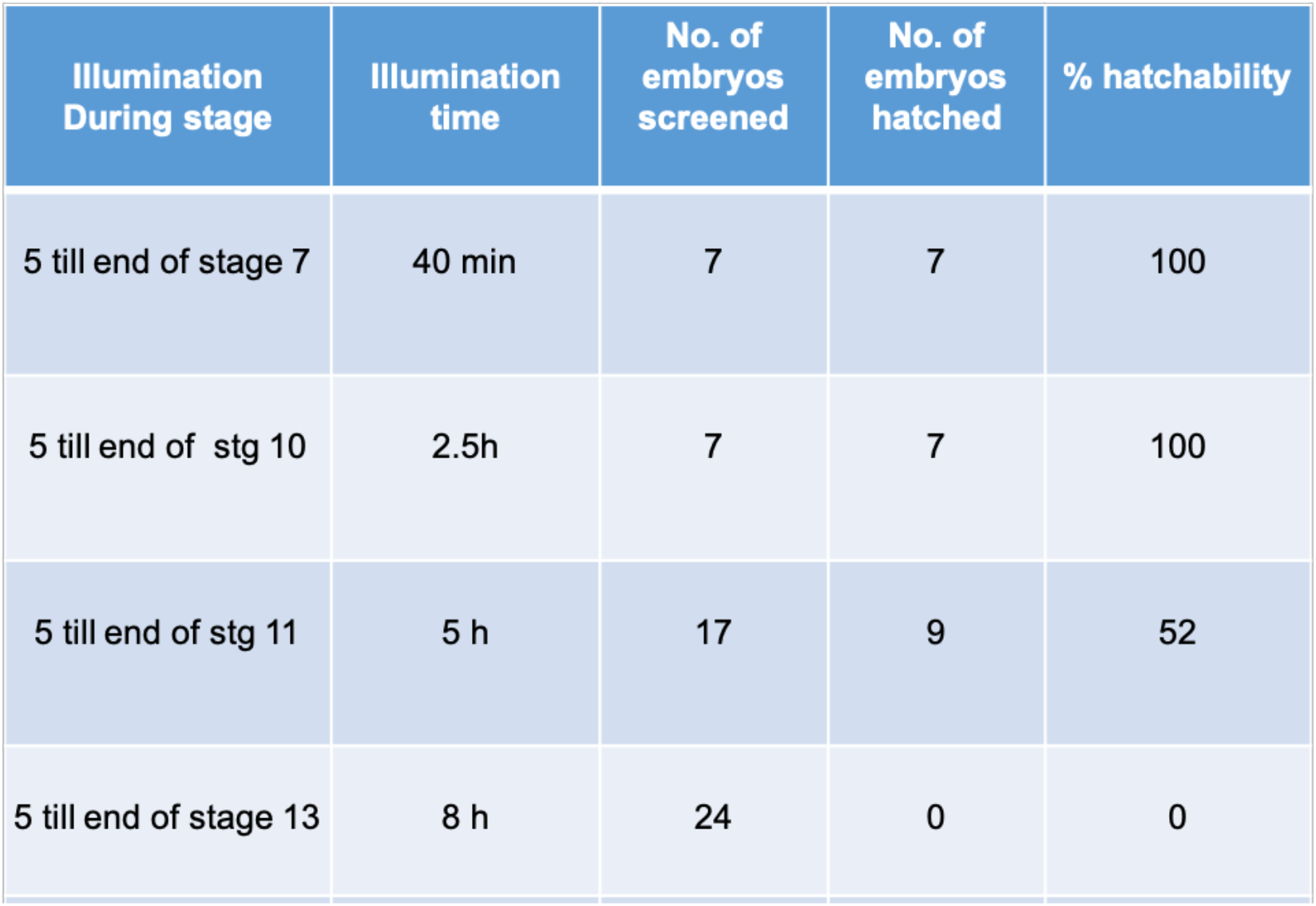
Hatchability of Twi-GAL4>Opto-htl embryos illuminated during different time windows.

| Illumination During stage | Illumination time | No. of embryos screened | No. of embryos hatched | % hatchability |
| --- | --- | --- | --- | --- |
| 5 till end of stage 7 | 40 min | 7 | 7 | 100 |
| 5 till end of stg 10 | 2.5h | 7 | 7 | 100 |
| 5 till end of stg 11 | 5 h | 17 | 9 | 52 |
| 5 till end of stage 13 | 8 h | 24 | 0 | 0 |

We observed that activation of Opto-htl during the developmental period from late stage 10 until late stage 12 was both necessary and sufficient to induce the described mesodermal defects. This window is crucial for heart morphogenesis and requires precise levels of FGFR activation to ensure proper cell fate specification and robust development (Reim & Frasch, 2010). In contrast, later developmental processes (post stage 13) appear to be able to tolerate a much higher range of Htl activity, possibly because cell-fate specification has already happened by this time and/or cells can buffer any increased Htl activity.

## DISCUSSION

The embryonic mesoderm gives rise to a diverse range of cell types that form the heart, visceral musculature, haemocytes and the fat body. Htl plays a crucial role in shaping the mesoderm by ensuring (i) uniform spreading of the mesoderm over the ectoderm and (ii) subsequent fate specification of different tissue precursors (Michelson *et al*., 1998; Wilson *et al.,2000;* Stathopoulos *et al*., 2004; Kadam *et al*., 2009). Here, we have demonstrated the use of an optogenetic tool (Opto-htl) to study spatiotemporal regulation of Htl signaling by controlling its level of activation *in vivo* along with precisely defined spatial and temporal windows. We found that the embryo is highly sensitive to Htl over-activation during the time window from late stage 10 till end of stage 12 (early stages of heart morphogenesis). Illumination from stage 13 onwards did not result in any remarkable defects (Fig. 6). Within the stage 10 to 12 window, the severity of the phenotype decreased with decreasing light intensity (Fig. 5A). Opto-htl provides a distinct advantage over previous studies of Htl over-activation, as it allows careful tuning of the strength of the pathway response. We can induce a stronger signaling dosage response in the mesoderm from stage 10 by increasing the light intensity of illumination.

There was no apparent increase in the total number of mesodermal cells upon light activation in Twi-Gal4>Opto-htl embryos observed at stage 11(Fig. S2). By stage 13, however, we see an increase in the number of Hand::GFP-positive cells in these embryos when exposed to light (Movie S5B-C and Fig. 4C, compare the non-illuminated and illuminated sides). We currently do not have a live reporter for cardioblasts with clear expression at stage 10-12 (Hand::GFP comes up in stage 13). In related work, we are developing new lines to explore the dynamics of early heart formation. We can then unravel whether the changes in cell number are related to over-proliferation of Tin-positive cardioblasts or switches in cell identity of other mesodermal cells and also decipher the exact timing of these events during development.

Surprisingly, illuminated Twi-Gal4>Opto-htl embryos at 1mW intensity do not show an increase in number of Eve-positive precursors (at stage 11) that form future pericardial cells and DA1 muscles. This contrasts with the htl-λ construct, that induced additional Eve-positive precursors at stage 11 but did not result in a lethal phenotype at later stages (Fig. S3). In Twi-Gal4>Opto-htl embryos, Eve-positive DA1 muscles at stage 16 are completely missing from light-activated (1mW) embryos at stage 16 (Fig. 3C). At low light activation, our Opto-htl phenotype was similar to that of htl-λ with respect to the number of ectopic Tin-positive heart cells observed at stage 16 (Fig. S3, asterisk). It is possible that at higher light intensity of 1mW, Opto-htl over-activation of the FGFR pathway is inducing conversion from a muscle cell fate to a cardiac or pericardial cell fate; *i.e*. Mef-2 positive muscle founders adopting a Tin-positive cardioblast fate and Eve-positive dorsal muscle cells adopting a pericardial fate. Therefore, Opto-htl potentially enables fine-tuning of FGFR-dependent cell fate decisions within the developing mesoderm. This opens up the possibility to explore the role of *htl* activity in controlling mesoderm cell behaviour with greater precision and across a greater range than possible with other tools.

There are several different pathways downstream of Htl (Ras/Raf/MAPK, PLCγ, PI3/AKT) characterised based on their molecular components and the response they generate (Ornitz & Itoh., 2015). Each pathway activates transcription of a set of different genes generating cellular responses such as proliferation, differentiation, migration, cell survival, and apoptosis. The MAPK pathway activation plays an important role in development of several different cancer types (Roskoski., 2019). We compared the developmental effect of constitutive activation at the receptor level with Opto-htl (potentially activating all downstream pathways) to activation of only the Ras/Raf/MAPK pathway in the mesoderm by driving Opto-SOS (Johnson *et al*., 2017) with Twi-Gal4. Upon illumination, we observe similar phenotypes in the heart as with Opto-htl; abundance of Tin-positive cardioblasts and loss of DA1 muscles (Fig. S4). The phenotypes are pronounced at 4.5mW illumination intensity, at which no DA1 muscles are observed and there is an abundance of Eve-positive pericardial cells and Tin-positive cardioblasts. At lower intensities, some of the DA1 structure is maintained. Hyperactivation of the Ras/Raf/MAPK pathway alone leads to phenotypes similar to Opto-htl in the mesoderm. This suggests that MAPK signalling dosage is critical in regulating cell proliferation and cell fate specification events in the mesoderm during stages 10-12.

We observed that Opto-htl can restore Tin-positive cardioblasts in *htl* null mutants upon light activation (Fig. 2C). Specification of Tin-positive heart precursors relies on the dorsal-most mesodermal cells receiving cues from Decapentaplegic and Wingless (Reim & Frasch., 2010), which emanate from the neighbouring ectodermal tissue. In *htl*-null mutants, mesoderm spreading is defective, but some cells still lie near to the neighbouring ectoderm. This likely explains how Opto-Htl is able to activate *tin* expression in *htl* null mutants and why the resulting pattern of Tin-positive cardioblasts is heterogeneous. This is consistent with the role of Htl in heart cell specification, independent from its role in mesoderm spreading (Michelson *et al*, 1998).

Our results suggest that Htl-dependent cell fate decisions are highly stage specific and dosage dependent. An increase in dosage leads to variations in different cell types of the heart and muscle lineage. These events are specifically restricted to a time window for Htl over-activation between late stage 10 and early stage 13, though recall that the lower time bound is uncertain due to relatively lower expression levels of Twi-Gal4>Opto-htl at early stages. Using Opto-htl to control several parameters of FGFR activation - dosage, spatial region of activation, and timing of activation - we have provided insight into different modes of Htl action in the developing mesoderm. In future work, we aim to fine tune these parameters further, to study how they affect cell fate changes *in htl* null embryos.

FGFR signalling plays a crucial role in the development and maintenance of several different organ systems in humans (heart, lungs, brain, skeletal muscles, etc.) and is also a target in disease therapies (Xie *et al*., 2020). There is evidence from animal studies that FGFR signal activation has potential for use in tissue regeneration and repair (Engel *et al*., 2006; Korf-Klingebiel *et al*., 2011; House *et al*., 2015). Optogenetic control over signalling pathways allows us to fine tune their activation or deletion in a highly controlled spatiotemporal manner. Studying the conditional effects of these manipulations on development and disease pathogenesis has the potential to lead to novel therapeutics.

## Acknowledgements

We thank Kenji Matsuno, Maria Leptin and Zhe Han for fly lines. We thank Manfred Frasch, James Sharpe and Eileen Furlong for the Tinman, Eve and Mef-2 antibodies respectively. We acknowledge members of the Saunders laboratory for fruitful discussions. This work was supported by a Singapore Ministry of Education AcRF Tier 3 Grant (MOE2016-T3-1-002).

## Author Contributions

All authors designed and planned the study. VY performed all experiments and data analysis, with guidance from NT and TES. VY and TES wrote the initial manuscript, with all authors contributing to the final manuscript.

## Materials and Methods

### Fly stocks, husbandry and genetics

UAS-htl-Cry2mCherry construct was generated using Gibson assembly (NEB) of four fragments: pPW vector backbone digested and purified, synthesized src42A myristoylation (myr) signal sequence, PCR amplified cytoplasmic region of htl/btl from cDNA, PCR amplified CRY2mCherry sequence from the AddGene plasmid #26866 (Kennedy *et al.,* 2010). The recombinant plasmid was sent to BestGene Inc for P-element transformation. Twi-Gal4 (BDSC58804, BDSC2517) was used to drive expression in the mesoderm and Byn-Gal4>UAS-myr-GFP was used to drive expression in the hindgut (kindly provided by Kenji Matsuno). To mark the heart cells for live imaging, we used Hand::GFP (GFP driven by the Hand cardiac and hematopoietic (HCH) enhancer (Han, 2006)) and formed a stable line: Hand::GFP;UAS-htl-Cry2mCherry.

For the rescue experiments, UAS-htl-Cry2mCherry on 2^nd^ chromosome was combined with the *htl^AB42^* null mutant on 3^rd^ chromosome (BDSC 5370). Similarly, Twi-Gal4 on 2^nd^ chromosome (BDSC2517) was combined with *htl^AB42^* null mutant to form a stable line. UAS-htl-Cry2mCherry; htl^AB42^ / TM3-ftz-lacz virgin females were crossed to Twi-Gal4; htl^AB42^ / TM3-ftz males resulting in 25% homozygous mutant embryos expressing Twi-Gal4>UAS-htl-Cry2mCherry. All fly lines were raised at 25°C.

### Identification of homozygous *htl* mutant embryos for rescue experiments

For the rescue crosses, a balancer chromosome carrying a lacZ-transgene was used. lacZ expression was driven by the ftz-promoter and was detected in embryos using anti-β-Gal antibody (DSHB 40-1a). For cross-section staining, lacZ-positive and lacZ-negative embryos were sorted under a dissecting microscope based on the β-gal staining pattern prior to cutting in 70% glycerol using a 25-gauge needle.

### Immunostaining

Embryos were collected at the desired stages, then dechorionated using bleach and fixed in heptane saturated with 37% paraformaldehyde (PFA) for 50 minutes. The vitelline membrane was subsequently removed using a needle. Prior to immunostaining, the embryos were blocked in 10% BSA-PBS. Antibodies used were mouse anti-β-Gal (1:100, DSHB), rabbit anti-mCherry (1:100, Abcam), guinea pig anti-Eve (1:800, kindly provided by James Sharpe), rabbit anti-dpERK (1:100, CST), mouse anti-fas3 (1:300, DSHB), Rabbit anti-tin (1:1000, kindly provided by Manfred Frasch), mouse anti-myosin heavy chain (1:100, DSHB), rabbit anti-mef2 (1:800, kindly provided by Eileen Furlong). Primary antibodies were detected with Alexa Fluor-labelled secondary antibodies (1:500; LifeTech). Embryos were mounted in AquaMount (PolySciences, Inc.) and imaged using a Zeiss LSM710 microscope with a C-Apochromat 40x water-immersion objective or a Nikon A1-RS scanning confocal microscope with 40x water-immersion objective. For dpERK intensity comparisons, both dark and light condition embryos were collected, stained and imaged under same conditions (antibody dilutions, laser power, etc).

### Illumination experiments

To score hatchability, embryos were imaged on a bright-field stereomicroscope at 25°C. To maintain the dark condition where needed, embryos were observed with the light source covered with amber paper to block 488nm light. For illuminating embryos, we used a Nikon LED light base as the light source and measured different light intensities using an intensity power meter set at the 488nm range. All experiments were carried out at 1 mW (measured at the sample plane) unless otherwise stated. For different time-window illuminations, stage 5 embryos were collected in the dark condition (488nm light blocked by amber paper) and then exposed to light. For keeping embryos in dark for recovery after illumination, we used a box covered with aluminium foil. For staining experiments, dark and light condition embryos were fixed and stained simultaneously at exactly similar stages. All embryos were collected and staged according to Ortega & Hartenstein (1985).

### Light-sheet microscopy for spatially controlled activation

Embryos were dechorionated using bleach, mounted into a capillary containing 1% low-melting agarose (Sigma) in an upright position and imaged on a Z.1 light-sheet fluorescence microscope (Carl Zeiss, Germany) using a 40x water immersion objective. The light-sheet microscope was equipped with 30 mW 488 nm and 20 mW 561 nm lasers with BP505–545 and LP585 emission filters respectively. Embryos were imaged using dual-side illumination by a light-sheet modulated into a pivot scan mode. The 488 nm excitation laser was used at 6% power with 7.5ms exposure time and the 561 nm excitation laser was used at 13% laser power with 20ms exposure time. For control embryos, only the 561 nm laser was used for scanning.

For spatial control of activation, embryos were mounted vertically and scanned with the 488nm laser from the lateral side such that the light-sheet was parallel to the plane of the embryo. ~35um of slices from the embryo surface were scanned so as to illuminate the heart precursors on only one side of the embryo. We scanned embryos with the 488nm laser from stage 10 till stage 13 and then we let embryos develop in the dark until stage 16 (no scanning with 488nm laser). At stage 16, we scanned the dorsal side of the embryos to image the heart and compared signals from the illuminated and non-illuminated sides.

---

**Supplementary Fig. 1.**
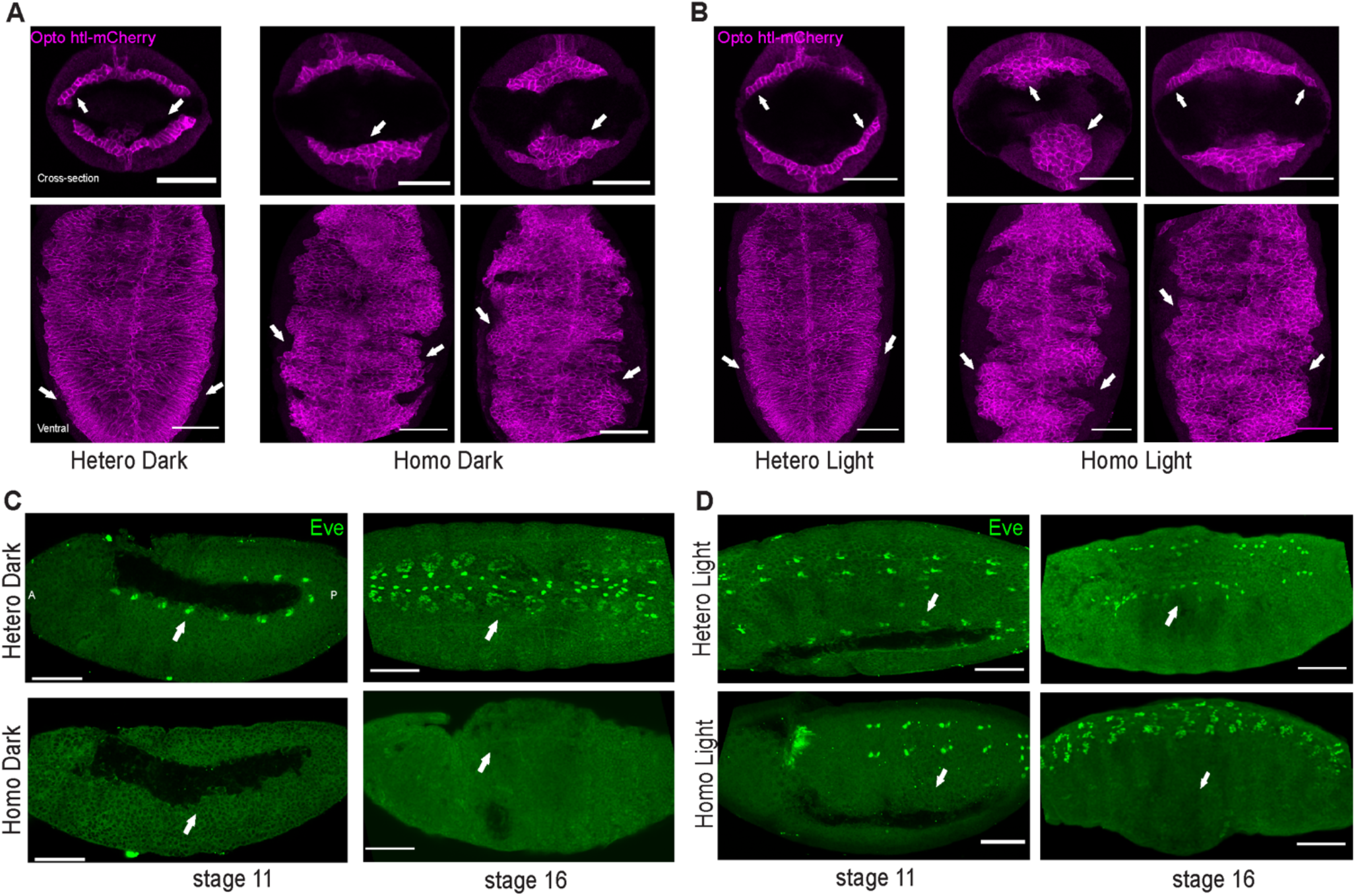
Rescue phenotypes using UAS-Opto-htl. A) Mesoderm spreading in heterozygous and homozygous *htl* mutant embryos kept under dark conditions, imaged at the cross-section (top row) and from the ventral side (bottom row). B) Mesoderm spreading in heterozygous and homozygous *htl* mutant embryos illuminated from stage 5 till stage 10. Upon illumination, homozygous mutants expressing Opto-htl exhibit clustering of mesoderm cells and non-uniform spreading with flattening observed occasionally in some regions. No rescue of Eve-positive cells is observed in the homozygous mutants under light either at stage 11 (C) or stage 16 (D). Scale bar = 50μm.

**Supplementary Fig. 2.**
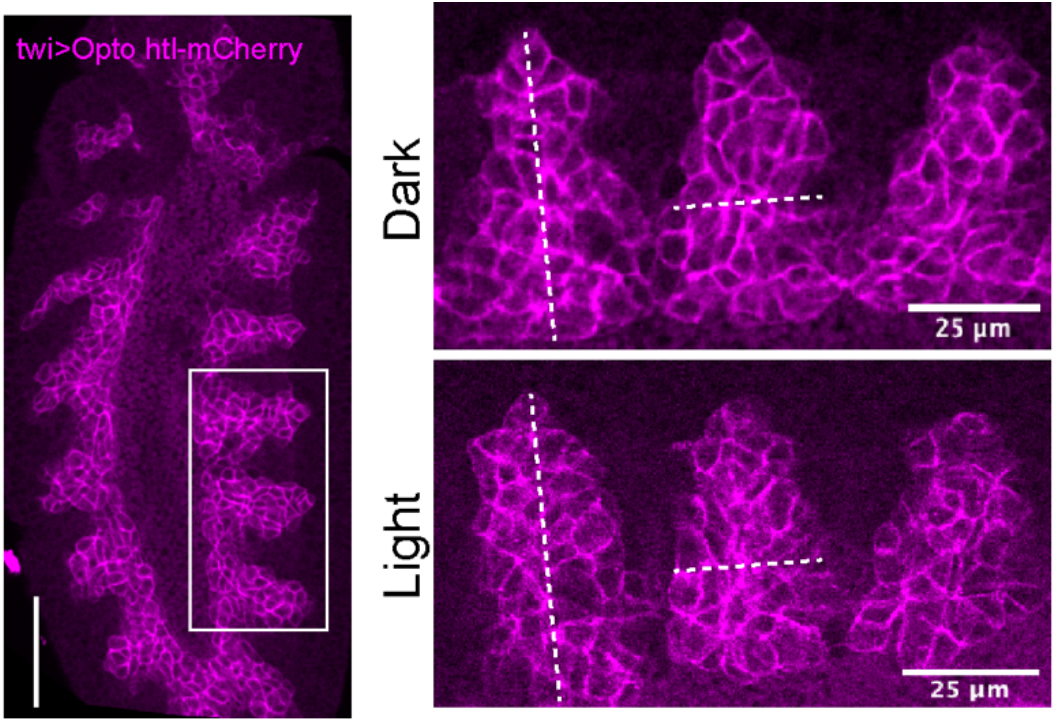
Mesodermal cells in Opto-htl embryos. Twi-Gal4>Opto-htl-mCherry embryos fixed and stained at stage 11 with anti-mCherry antibody under dark and light conditions. No expansion of mesodermal cells is observed upon light activation. Scale bar = 50μm unless stated otherwise.

**Supplementary Fig. 3.**
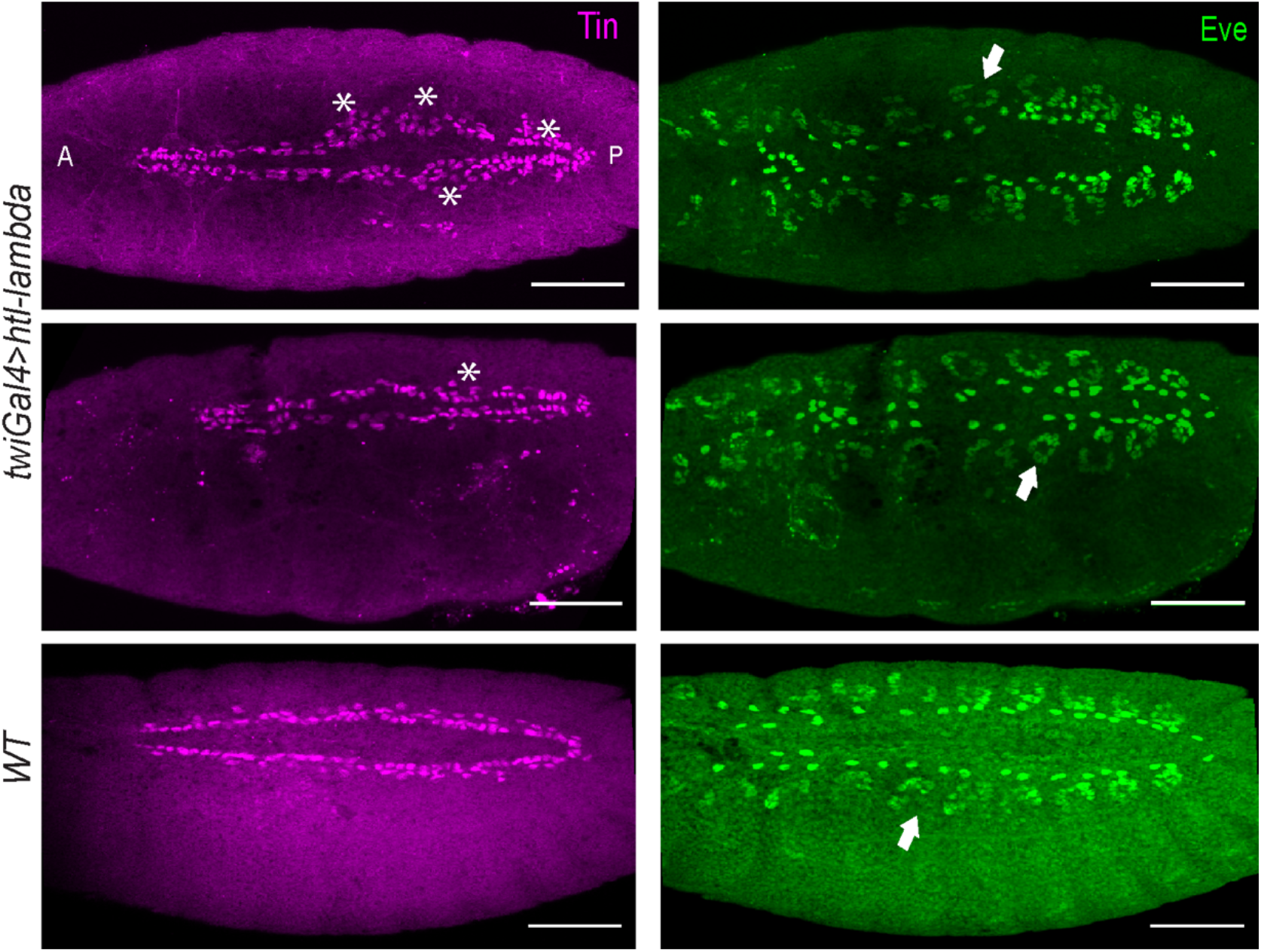
htl-λ activation in the mesoderm. Dorsal view of Twi-Gal4>UAS-htl-λ embryos fixed and stained at stage 15 with Tin and Eve antibody, compared with WT embryos at the same stage. Asterisks represent ectopic tin cells. Arrowhead point towards eve-positive DA1 muscles and pericardial cells in a given hemi segment. Scale bar = 50μm.

**Supplementary Fig. 4.**
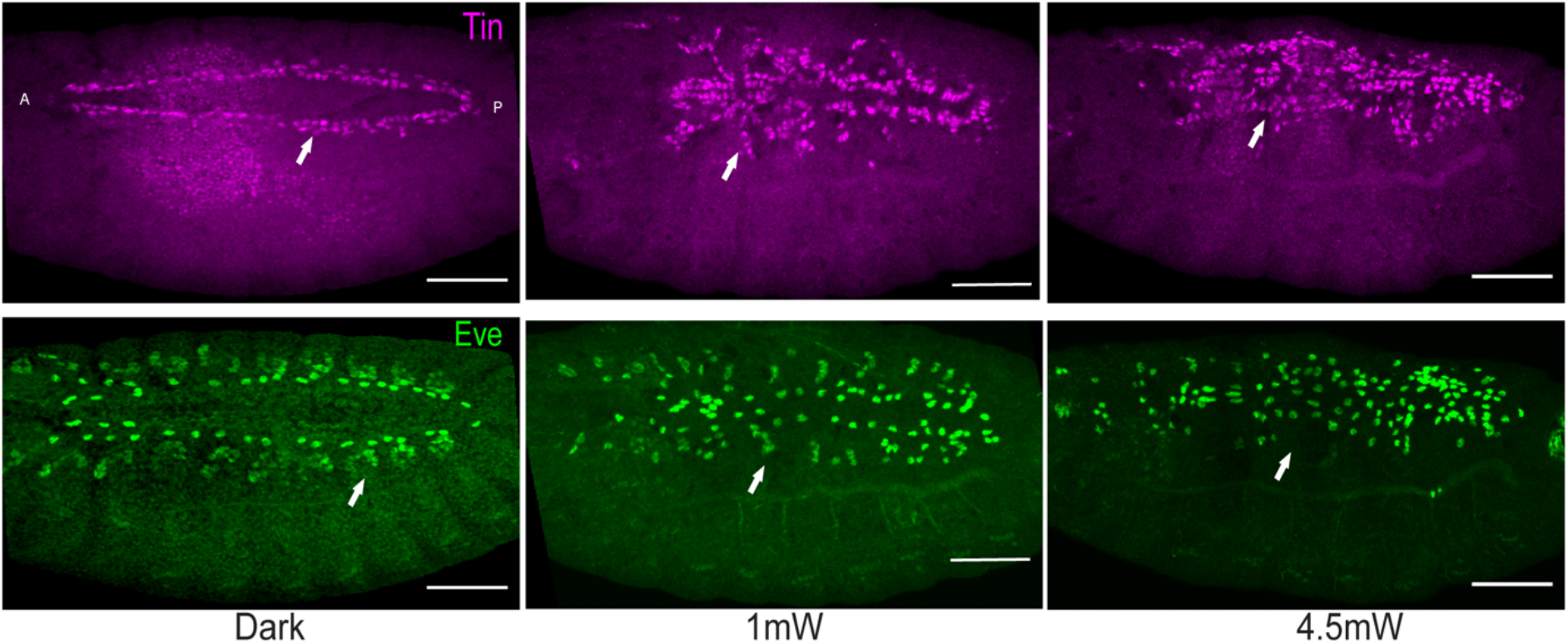
Opto-SOS activation in the mesoderm. Dorsal view of Twi-Gal4>UAS-Opto-SOS embryos fixed and stained at stage 16 with Tin and Eve antibody. Embryos kept under dark show normal heart development with Tin and Eve patterns similar to wild-type. Embryos illuminated at 1mW and 4.5mW light intensity from stage 5 till stage 16 show a progressive increase in both Tin and Eve positive cells with a complete loss of the DA1 muscle group (arrowheads) at 4.5mW intensity. Scale bar = 50μm.

## Movie Legends

**Supplementary Movie 1: Opto-htl expression in the mesoderm during embryogenesis**

Twi-Gal4>Opto-htl embryo scanned with 561nm laser at 3 min intervals from stage 5 until end of embryogenesis to visualise expression of Opto-htl.

**Supplementary Movie 2: Hatchability score for Opto-htl embryos under dark conditions**

Twi-Gal4>Opto-htl embryos imaged on a stereoscope covered with amber paper (to block 488nm light) from stage 5 until hatching. (Dark condition).

**Supplementary Movie 3: Hatchability score for Opto-htl embryos under constant illumination**

Twi-Gal4>Opto-htl embryos imaged on a stereoscope from stage 5 until end of embryogenesis using a 488nm light intensity of 1mW measured at the sample plane. (Light condition).

**Supplementary Movie 4: Cardioblast migration and matching in Opto-htl embryo kept under dark conditions**

Twi-Gal4>hand::GFP; Opto-htl embryo kept under dark condition until stage 13 and then imaged to visualise the migration of the GFP positive heart cells until matching at stage 16.

**Supplementary Movie 5: Cardioblast migration and matching in Opto-htl embryo illuminated with 488nm laser**

Twi-Gal4>hand::GFP;Opto-htl embryos illuminated with 488nm laser from late stage 10 until early stage 13 and then imaged using a light-sheet microscope to visualise the migration of the GFP positive heart cells until matching at stage 16. (a) represents dorsal view; (b) and (c) represent lateral views of the same embryo.

## Sequences and Primers

### Myr signal sequence (src42A)

ATGGGTAACTGCCTCACCACACAGAAGGGCGAACCCGACAAGCCCGCA

### Htl FGF cytoplasmic region (Amino acid position 331-729)

(Ref. http://www.uniprot.org/uniprot/Q07407)

TATGCCATCCGAAAGATGAAACATGAAAAGGTGTTGAAACAACGCATCGAAACCGTTCACCAGTGGACCAAGAAAGTGAT CATCTTCAAGCCCGAAGGTGGCGGAGACTCCAGTGGTTCCATGGACACCATGATTATGCCGGTGGTTAGGATACAGAAAC AGCGCACCACTGTTCTTCAGAATGGCAACGAGCCGGCTCCATTCAATGAATATGAATTTCCACTGGACTCGAACTGGGAA CTGCCCAGAAGTCATTTGGTACTGGGTGCCACTTTGGGAGAAGGTGCTTTCGGACGAGTGGTCATGGCGGAGGTCAATAA TGCCATTGTCGCCGTGAAAATGGTGAAGGAAGGACACACGGATGATGACATTGCCAGCTTGGTGCGGGAAATGGAAGTGA TGAAGATCATTGGGCGACATATCAATATTATTAACTTACTTGGTTGCTGCAGTCAAAATGGTCCGCTCTATGTGATTGTC GAGTATGCGCCACACGGAAATCTCAAGGACTTCCTCTATAAAAATCGACCCTTCGGAAGGGATCAGGATAGGGACAGCTC GCAACCGCCGCCATCGCCACCAGCTCATGTGATAACCGAAAAGGATCTGATCAAATTTGCCCACCAAATTGCCAGAGGAA TGGACTATTTGGCCTCGCGGCGATGCATCCATCGAGATTTGGCAGCCAGGAATGTGCTCGTCAGCGATGATTATGTGCTG AAGATTGCTGATTTTGGACTGGCGAGGGACATTCAAAGCACGGATTACTATCGGAAGAACACAAATGGCAGGCTACCCAT CAAATGGATGGCACCGGAGTCGCTGCAGGAGAAATTCTATGATTCCAAGAGCGATGTCTGGTCATATGGCATCCTGCTGT GGGAGATCATGACCTATGGGCAGCAACCATATCCAACTATCATGTCCGCTGAGGAGCTGTACACCTATCTCATGTCCGGT CAGCGGATGGAGAAACCAGCGAAATGCTCCATGAACATCTACATTCTGATGCGACAATGTTGGCATTTCAACGCCGACGA TCGGCCACCTTTTACGGAAATTGTTGAGTATATGGACAAGCTGCTCCAGACGAAGGAGGACTACCTCGATGTGGATATCG CCAATCTGGATACACCGCCCTCGACTAGCGACGAGGAGGAAGATGAAACGGACAACCTGCAGAAGTGGTGTAATTAT

### CRY2mcherry (Ref. Addgene26866)

atgaagatggacaaaaagactatagtttggtttagaagagacctaaggattgaggataatcctgcattagcagcagctgc tcacgaaggatctgtttttcctgtcttcatttggtgtcctgaagaagaaggacagttttatcctggaagagcttcaagat ggtggatgaaacaatcacttgctcacttatctcaatccttgaaggctcttggatctgacctcactttaatcaaaacccac aacacgatttcagcgatcttggattgtatccgcgttaccggtgctacaaaagtcgtctttaaccacctctatgatcctgt ttcgttagttcgggaccataccgtaaaggagaagctggtggaacgtgggatctctgtgcaaagctacaatggagatctat tgtatgaaccgtgggagatatactgcgaaaagggcaaaccttttacgagtttcaattcttactggaagaaatgcttagat atgtcgattgaatccgttatgcttcctcctccttggcggttgatgccaataactgcagcggctgaagcgatttgggcgtg ttcgattgaagaactagggctggagaatgaggccgagaaaccgagcaatgcgttgttaactagagcttggtctccaggat ggagcaatgctgataagttactaaatgagttcatcgagaagcagttgatagattatgcaaagaacagcaagaaagttgtt gggaattctacttcactactttctccgtatctccatttcggggaaataagcgtcagacacgttttccagtgtgcccggat gaaacaaattatatgggcaagagataagaacagtgaaggagaagaaagtgcagatctttttcttaggggaatcggtttaa gagagtattctcggtatatatgtttcaacttcccgtttactcacgagcaatcgttgttgagtcatcttcggtttttccct tgggatgctgatgttgataagttcaaggcctggagacaaggcaggaccggttatccgttggtggatgccggaatgagaga gctttgggctaccggatggatgcataacagaataagagtgattgtttcaagctttgctgtgaagtttcttctccttccat ggaaatggggaatgaagtatttctgggatacacttttggatgctgatttggaatgtgacatccttggctggcagtatatc tctgggagtatccccgatggccacgagcttgatcgcttggacaatcccgcgttacaaggcgccaaatatgacccagaagg tgagtacataaggcaatggcttcccgagcttgcgagattgccaactgaatggatccatcatccatgggacgctcctttaa ccgtactcaaagcttctggtgtggaactcggaacaaactatgcgaaacccattgtagacatcgacacagctcgtgagcta ctagctaaagctatttcaagaacccgtgaagcacagatcatgatcggagcagcaGCCCGGGATCCACCGGTCGCCACCAT GGTGAGCAAGGGCGAGGAGGATAACATGGCCATCATCAAGGAGTTCATGCGCTTCAAGGTGCACATGGAGGGCTCCGTGA ACGGCCACGAGTTCGAGATCGAGGGCGAGGGCGAGGGCCGCCCCTACGAGGGCACCCAGACCGCCAAGCTGAAGGTGACC AAGGGTGGCCCCCTGCCCTTCGCCTGGGACATCCTGTCCCCTCAGTTCATGTACGGCTCCAAGGCCTACGTGAAGCACCC CGCCGACATCCCCGACTACTTGAAGCTGTCCTTCCCCGAGGGCTTCAAGTGGGAGCGCGTGATGAACTTCGAGGACGGCG GCGTGGTGACCGTGACCCAGGACTCCTCCCTGCAGGACGGCGAGTTCATCTACAAGGTGAAGCTGCGCGGCACCAACTTC CCCTCCGACGGCCCCGTAATGCAGAAGAAGACCATGGGCTGGGAGGCCTCCTCCGAGCGGATGTACCCCGAGGACGGCGC CCTGAAGGGCGAGATCAAGCAGAGGCTGAAGCTGAAGGACGGCGGCCACTACGACGCTGAGGTCAAGACCACCTACAAGG CCAAGAAGCCCGTGCAGCTGCCCGGCGCCTACAACGTCAACATCAAGTTGGACATCACCTCCCACAACGAGGACTACACC ATCGTGGAACAGTACGAACGCGCCGAGGGCCGCCACTCCACCGGCGGCATGGACGAGCTGTACAAGTAA

### Primers for amplifying FGF cytoplasmic region

FP 5′ – GAACCCGACAAGCCCGCATATGCCATCCGAAAGATGAAAC –3′

RP 5′ – ctttttgtccatcttcatATAATTACACCACTTCTGCAGGTTGTC–3′

### Primers for amplifying CRY2-mcherry

FP– 5′–CAGAAGTGGTGTAATTATatgaagatggacaaaaagactatagtttgg–3′

RP 5′– GGGGTGCCTAATGCGGCCGCTTACTTGTACAGCTCGTCCATGC –3′

### Myr sequence synthesized

5′–GGTATACACCTAGGCGGTACCATGGGTAACTGCCTCACCACACAGAAGGGCGAACCCGACAAGCCCGCATATGCCATCCGAAAGATG–3′

3′–CCATATGTGGATCCGCCATGGTACCCATTGACGGAGTGGTGTGTCTTCCCGCTTGGGCTGTTCGGGCGTATACGGTAGGCTTTCTAC–5′

